# Integrating exposure knowledge and serum suspect screening as a new approach to biomonitoring: An application in firefighters and office workers

**DOI:** 10.1101/630848

**Authors:** Rachel Grashow, Vincent Bessonneau, Roy R. Gerona, Aolin Wang, Jessica Trowbridge, Thomas Lin, Heather Buren, Ruthann A. Rudel, Rachel Morello-Frosch

## Abstract

**Background:** Women firefighters are exposed to recognized and probable carcinogens, yet there are few studies of chemical exposures and associated health concerns, such as breast cancer. Biomonitoring often requires *a priori* selection of compounds to be measured, and so may not detect important, lesser known, exposures.

**Objectives:** The Women Firefighters Biomonitoring Collaborative (WFBC) created a biological sample archive and conducted a general suspect screen (GSS) to address this data gap.

**Methods:** Using liquid chromatography-quadrupole time-of-flight mass spectrometry (LC-QTOF/MS) we sought to identify candidate chemicals of interest in serum samples from 83 women firefighters (FF) and 79 office workers (OW) in San Francisco. Through the GSS approach we identified chemical peaks by matching accurate mass from serum samples against a custom chemical database of 740 slightly polar phenolic and acidic compounds, including many of relevance to firefighting or breast cancer etiology. We then selected chemicals for confirmation based on *a priori* criteria: 1) detection frequency or peak area differences between OW and FF; 2) evidence of mammary carcinogenicity, estrogenicity, or genotoxicity; and 3) not currently measured in large biomonitoring studies.

**Results:** We detected 620 chemicals that matched 300 molecular formulas in the WFBC database, including phthalate metabolites, phosphate flame retardant metabolites, phenols, pesticides, nitro-and nitroso-compounds, and per- and polyfluoroalkyl substances. The average number of chemicals from the database that were detected in participants was 72 and 70 in FF and OW, respectively. We confirmed 8 of the 20 prioritized suspect chemicals –including two alkylphenols, ethyl paraben, BPF, PFOSAA, benzophenone-3, benzyl p-hydroxybenzoate, and triphenyl phosphate--by running a matrix spike of the reference standards and using m/z, retention time and the confirmation of at least two fragment ions as criteria for matching.

**Conclusion:** GSS provides a powerful high-throughput approach to identify and prioritize novel chemicals for biomonitoring and health studies.

## INTRODUCTION

Firefighters are exposed to complex and variable chemical mixtures that include known carcinogens. In addition to exposures during fire suppression activities (Adetona et al. 2013; Fent et al. 2014, 2018; Navarro et al. 2017; Pleil et al. 2014), firefighters pick up chemical exposures from their equipment, such as fire extinguishing foams or protective gear (Alexander and Baxter 2016; Fent et al. 2015), and also from automotive diesel (Oliveira et al. 2017). These compounds include benzene, polycyclic aromatic hydrocarbons (PAHs), nitro-PAHs, formaldehyde, dioxins, flame retardants, polychlorinated biphenyls, and poly- and perfluorinated substances (PFAS) (Caux et al. 2002; Feunekes et al. 1997; Moen and Ovrebø 1997; Waldman et al. 2016). These chemicals are associated with a wide range of cancers and other health effects in human and experimental animal studies, and it is noteworthy that many of these exposures have been identified as potential breast carcinogens either because they cause mammary gland tumors in laboratory animals, or because they alter mammary gland development (Rudel et al. 2011, 2014).

Research examining the chemical exposures and health risks faced by firefighters, and women firefighters in particular, is limited. A 2015 study conducted by the National Institute for Occupational Safety and Health (NIOSH) on 19,309 male US firefighters observed positive associations between the total time spent at fires and lung cancer incidence and mortality, and between the total number of response to fires and leukemia mortality from 1950-2009 (Daniels et al. 2015). An earlier report from this NIOSH cohort that included 991 women showed non-significant increases in breast cancer incidence and mortality in both men and women, compared with the general US population; these increases were largest at younger ages (<65 for men, 50-55 for women) (Daniels et al. 2014). Studies in multiple countries have also documented an elevated risk of certain cancers in male firefighters and other first responders, including thyroid, bladder, kidney, prostate, testicular, breast, brain, digestive cancers, multiple myeloma, and non-Hodgkin’s lymphoma (Ahn et al. 2012; Bates 2007; Delahunt et al. 1995; Kang et al. 2008; Ma et al. 2005, 2006; Tsai et al. 2015). A meta-analysis of 32 studies determined an increased risk of certain cancers in the mostly male firefighter population (LeMasters et al. 2006). Most studies do not calculate risks to female firefighters; however, in a study on cancer incidence among Florida professional firefighters, female firefighters showed a significantly increased risk of cancer overall, as well as Hodgkin’s lymphoma disease and thyroid cancer, compared with the Florida general population (Ma et al. 2006). Although women make up 5.1% of firefighters across the United States, (US Department of Labor 2018) their numbers can be higher in urban jurisdictions, including in San Francisco, which has one of the highest proportions of women firefighters (15%) (Hulett et al. 2008). As fire departments diversify and increase the number of women in their ranks, it is important to characterize chemical exposures and implications for health outcomes of particular relevance to women, such as breast cancer, that might not be addressed in existing studies, which have been primarily conducted among men.

Biomonitoring is an important tool in environmental and occupational health studies seeking to link health outcomes to chemical exposures. External measurements including in air, dust, and water do not always reflect internal dose, and biomonitoring studies in human tissue can integrate over multiple routes of exposure including dermal, inhalation and ingestion. One limitation of many biomonitoring studies is that they rely on *a priori* selection of targeted chemicals for study. This *a priori* selection approach often lacks critical information about which chemicals are present in occupational settings (Egeghy et al. 2012; Judson et al. 2009), and about metabolic transformations. As a result, significant time and resources may be expended to develop analytical methods to measure chemicals without knowing whether they are present in biological specimens. For example, 20% of the 250 chemicals biomonitored in NHANES since 1999 were not detected in 95% or more of the US population, indicating that the criteria for selecting chemicals for biomonitoring has not always identified chemicals with prevalent exposure (CDC 2009). A more efficient and systematic approach is needed to identify a broader spectrum of environmental chemicals present in the human body; this strategy is now recognized as a critical component of an “exposome” approach (Buck Louis et al. 2013; Rappaport 2011; Wild 2012). One way to characterize the human exposome is to perform a general suspect screen (GSS) of biospecimens using high-resolution mass spectrometry. Recent applications of this approach identified novel chemical exposures among pregnant women, including benzophenone-1 and bisphenol S (Gerona et al. 2018; Wang et al. 2018).

To better understand how women firefighters are exposed to potential breast carcinogens and other understudied chemicals, we undertook a community-based, participatory biomonitoring project, a partnership among firefighters, environmental health scientists and environmental health advocates, known as the Women Firefighters Biomonitoring Collaborative, to develop a biospecimen archive of women firefighters and office workers in San Francisco. As part of the WFBC, we conducted a cross-sectional chemical biomonitoring study to identify novel chemical exposures by applying a discovery-driven, general suspect screen (GSS) using high-resolution mass spectrometry. Our goal was to characterize multiple chemical exposures, assess whether these exposures differ between firefighters and office workers, and prioritize candidate compounds for confirmation and targeted methods development. Ultimately, we applied a GSS approach to advance discovery of novel environmental chemicals in human biomonitoring.

## METHODS

### Study design

The Women Firefighter Biomonitoring Collaborative (WFBC) was designed to measure and compare exposures to potential breast carcinogens and other endocrine disrupting compounds (EDCs) in two occupational cohorts--women firefighters (FF) and office workers (OW) from the City of San Francisco, California, and to create an archive of biological specimens for exposomics research. The GSS was performed on serum samples collected from female firefighters and office workers using liquid chromatography-quadrupole time-of-flight mass spectrometry (LC-QTOF/MS) to characterize a wide spectrum of exposures to candidate compounds in our study population. This method screens for hundreds of acidic or phenolic organic compounds of interest, so the results represent a significantly larger universe of compounds in a biospecimen rather than a limited set of chemicals selected, *a priori*, for quantification. Accurate mass of each unique molecule (i.e. mass-to-charge ratio, *m/z*) generated by the LC-QTOF/MS was matched to chemical formulas from a custom database of 740 chemicals of interest, based on their relevance to firefighting and breast cancer etiology. From this WFBC database, we compared detection frequencies and peak areas of candidate compounds between firefighters and office workers to identify those that might be work-related. We then systematically combined expert knowledge on the sources, uses and toxicity of candidate compounds to prioritize and select a subset of chemicals for confirmation. Ultimately, we sought to demonstrate how GSS methods can be used to improve efficiencies in human biomonitoring by broadening the spectrum of potential environmental chemical exposures and applying exposure science expertise to identify and prioritize specific chemicals for confirmation by targeted analysis.

### Recruitment and consent

Women were eligible to participate in the WFBC study if they were over 18 years old, non-smokers, and employees of the City and County of San Francisco (office workers) or the San Francisco Fire Department (firefighters). In addition, firefighters had to have been working active duty for at least five years with the Department. Firefighters were recruited through letters, emails, and phone calls that targeted firefighter organizations, including United Fire Service Women, Local 798 of the International Association of Firefighters (IAFF), the Black Firefighters Association, Asian Firefighters Association, and Los Bomberos (Latino Firefighter Association). Informational meetings were held at the San Francisco Fire Department. Female office employees with the City and County of San Francisco were recruited through informational meetings, direct email, letters, telephone calls and by networking efforts through SEIU Local 1021. The study was publicized through regular newsletters and other online communication outlets regularly sent to firefighters and other San Francisco City and County employees through the Health Services System. WFBC study protocols were approved by the Institutional Review Board of the University of California, Berkeley (Protocol # 2013-07-5512). Informed consent was obtained prior to all interviews and sample collections. Subjects were not paid for participation, but did receive a $20.00 gift card and reimbursement to offset the cost of parking and transportation. Blood samples were collected between June 2014 and March 2015.

### Interviews and sample collection

Once consented and enrolled, participants were scheduled for an in-person interview and blood collection. Subjects met with a member of the research team to answer questions about their diet, home, job, other activities, and education. After completing the exposure interview, a trained phlebotomist drew blood samples, which were collected in four 10 mL red-top tubes without additives. Samples were collected at sites near participants’ work site and transported in a cooler with ice for processing within 3 hours of collection. Serum was separated by allowing it to clot at room temperature, then centrifuging at 3000 rpm for 10 minutes and −4°C. Serum was aliquoted into 1.2 mL cryo-vial tubes and stored at −80°C until analysis. All samples were processed and analyzed at the University of California, San Francisco. We collected and processed samples from 86 firefighters and 84 office workers. We analyzed serum samples from those who had sufficient serum for the chemicals analysis: from 83 firefighters and 79 office worker participants.

### WFBC suspect chemical database

To build a chemical database for our general suspect screen, we began with a database of 696 chemicals developed previously to identify environmental organic acids (EOA) among pregnant women, including chemicals from the following classes: phenols, such as parabens; phenolic and acidic pesticides and their predicted acidic and phenolic metabolites; per- and polyfluoroalkyl substances (PFAS); phthalate metabolites; phenolic metabolites of polybrominated diphenyl ethers (OH-BDEs) and polychlorinated biphenyls (OH-PCBs) (Wang et al. 2018). These EOAs include many common consumer product chemicals and environmental pollutants, as well as 356 predicted metabolites of common pesticides (Wang et al. 2018). We extended this EOA database for our WFBC analysis by adding environmental chemicals that were relevant to occupational exposures faced by firefighters and office workers and also chemicals implicated in breast carcinogenesis based on toxicological evidence. Specifically, we assessed the viability of adding over 100 chemicals, based on the following criteria: 1) chemicals shown to be rodent mammary gland carcinogens or that affect mammary gland development and so may increase breast cancer risk (Rudel et al. 2011, 2014); or 2) chemicals related to firefighting that could lead to occupational exposures, including perfluorinated compounds found in firefighting foams, and other flame retardants and their metabolites (Dodson et al. 2012, 2014; Rodgers et al. 2018). Chemicals that fit these two criteria were added to the WFBC database if their structures were expected to be compatible with the LC-QTOF/MS operating in negative ionization mode. For example, carcinogenic PAHs were not added to the database because they are unlikely be detected using this method. We were able to add 44 chemicals for a total of 740 in the WFBC database (Table S1).

### General suspect screening analysis using liquid-chromatography and quadrupole time-of-flight mass spectrometry (LC-QTOF/MS)

General suspect screening of serum was performed as previously described (Gerona et al. 2018). Briefly, 250 μL of serum was spiked with 2.5 μL of 1 mg/mL of internal standard (2.5 ng BPA-d16) and centrifuged at 3,000 rpm for 10 min. Analytes were extracted using solid-phase extraction (SPE; Waters Oasis HLB 10 mg, 1cc). Extracts were dried under a stream of nitrogen gas and reconstituted in 250 μL of 10% methanol.

Extracts were analyzed on a LC-QTOF/MS system consisting of an LC 1260 and a QTOF/MS 6550 (Agilent, Santa Cruz, CA, USA). Analytes were separated by reversed-phase chromatography using a C18 column (Agilent Poroshell 120, 2.1 mm × 100 mm, 2.7 mm particle size) maintained at 55°C. Mobile phase A consisted of water with 0.05% ammonium acetate (pH=7.8) and mobile phase B consisted of methanol with 0.05% ammonium acetate (pH=7.8). The elution gradient employed was: 0-0.5 min, 5% B; 1.5 min, 30% B; 4.5 min, 70% B; 7.5-10 min, 100% B; 10.01-14 min, 5% B. The injection volume was 50 μL.

Analyses were performed with a QTOF/MS operating in negative electrospray ionization mode (ESI-). Ions were collected in the *m/z* 80–600 range at high resolution for eluates coming out of the LC from 1-12 min. Using the Auto MS/MS mode (information-dependent acquisition), a product ion scan (MS/MS) of the three most abundant peaks at high resolution was triggered each time a precursor ion with an intensity of ≥500 counts/second was generated in the QTOF/MS scan using a collision voltage ranging from 0 to 40 V depending on ions *m/z*. The LC-QTOF/MS analysis produces a total ion chromatogram for each sample, which includes the following: the accurate mass of each unique compound (expressed as m/z of their respective anion), peak area, retention time (RT) and spectral data on the parent and fragment ions, including isotopic pattern.

We used the Agilent MassHunter Qualitative Analysis software Find-by-formula (FBF) algorithm to analyze QTOF/MS data for novel chemical exposures among firefighters and office workers using a set of optimized parameters previously reported (Gerona et al. 2018). First, all detected *m/z* were matched to potential compound hits in the WFBC chemical database. The algorithm imports molecular formulas from the database, automatically calculates their *m/z* values and then matches them to *m/z* measured by the QTOF/MS with a mass tolerance value of 10 ppm. A list of possible chemical matches was generated for all serum samples, which included the accurate mass (*m/z*), mass error (i.e. the difference between the experimental and the theoretical *m/z*), retention time (RT), peak area, and match scores (Schymanski et al. 2014). The initial LC-QTOF full scan identification resulted in 12,051 features with unique retention times, which matched to 300 chemical formulas in our WFBC database with multiple RTs/formula, or 620 unique chemical formula/RT combinations.

### Retention time correction and isomer distinction

Isomers (compounds with the same chemical formula but with different chemical structures) are recognized by the LC-QTOF method as the presence of multiple RTs, (measured in minutes) per chemical formula or mass. We distinguished isomers by clustering compounds based on RT. Briefly, we first ranked all suspect detections by RT for each chemical formula. We considered a suspect peak to be from a different isomer if its RT differed from the RT of the same chemical formula in the previous row by more than 0.16 minutes. Cutoff points ranging from 0.15 to 0.20 with a 0.01 increment were tested, and 0.16 allowed the best distinction based on graphical examination (Wang et al. 2018). Then, we aligned peaks originating from the same isomer to an identical RT. The final analytical sample consisted of 4,791 suspect detections that matched to 620 suspect chemicals (i.e., unique combinations of chemical formula and retention time).

### Chemical selection for validation and confirmation

We used a multi-step procedure and criteria to reduce the initial set of candidate chemical matches from the LC-QTOF/MS to a smaller set of compounds for validation by prioritizing matches that showed differences in exposure between firefighters and office workers or had toxicity characteristics relevant to breast cancer. We focused our general suspect screen on compounds in our database that were not pharmaceutical chemicals or chemicals that we had already identified for targeted analysis. We then used the following initial criteria to prioritize matches for validation: 1) at least 10% detection frequency difference between firefighters and office workers; 2) a higher peak area (indicator of higher relative concentration) in firefighters compared to office workers (paired t-test, p≤0.1); 3) ubiquitous chemicals detected in more than 90% of both firefighter and office worker groups and 4) whether a chemical had been flagged as a mammary carcinogen or mammary gland developmental disruptor [in (Rudel et al. 2007, 2011)]. As shown in Figure 2, this process yielded an initial list of 71 chemicals that we then narrowed down to 54 for potential confirmation based on the availability of an analytical standard.

In a second step for prioritizing tentative chemical matches for validation, we scored the remaining 54 chemicals based on the first set of selection criteria as well as the following additional characteristics: flame retardant chemicals, chemicals identified as estrogenic or genotoxic, chemicals not detected in office workers, and chemicals not currently biomonitored in NHANES (CDC 2019) or the California Biomonitoring Program (Biomonitoring California 2019) The specific criteria were chemicals: 1) listed as flame retardants [in (Dodson et al. 2012, 2014)]; 2) not detected in the office workers; 3) currently not biomonitored in NHANES or Biomonitoring California; 4) listed as “active” for at least one genotoxicity bioassay tested in PubChem (Wang et al. 2017); 5) listed as “active” for at least one estrogen receptor bioassay in PubChem (The PubChem Project). For bioassay data, results were downloaded from the PubChem website for each chemical. Then assay descriptions were queried for terms including “genotox*”, “estrogen” and “salmonella” (to flag all Ames assays). All assays matching those terms listed as “active” were tallied and chemicals with active assays were prioritized.

We scored the chemicals by assigning one point for each of the nine criteria. The study team reviewed the top scoring chemicals and selected twenty for validation based on score as well as data on uses, toxicity and sources using the Comparative Toxicogenomics Database (CTDB) (Davis et al. 2017), PubChem (Wang et al. 2017), Toxnet (Fowler and Schnall 2014), and the Toxin and Toxin Target Database (T3DB) (Wishart et al. 2015) (Table S2). Peaks that matched predicted pesticide metabolites in our database were not considered for validation because of the additional uncertainty about their presence in biological samples and lack of available reference standards.

### Confirmation of selected chemicals

We confirmed the presence of suspect chemicals in the serum samples by running the LC-QTOF/MS analysis using the corresponding reference standard spiked into synthetic serum. Tentative chemical matches from participant samples were confirmed if the *m/*z, at least two fragment peaks in the MS/MS spectra, and retention time of the authentic standard matched those found in the serum samples, consistent with level 1 confidence in identification (Schymanski et al. 2014).

### Statistical analysis

For statistical comparisons across demographic and occupational groups, we used the Wilcoxon rank sum test to compare continuous variables or the Fisher test for categorical variables. All data analysis and visualizations were completed using R, version 3.3.2 (R Core Team 2015).

## RESULTS

Table 1 shows the demographic characteristics for the 83 firefighters and the 79 office workers recruited for the WFBC study. At the time of recruitment, the San Francisco Fire Department (SFFD) had 224 active duty women firefighters who made up nearly 15% of its workforce. Among our study population, the average age of women firefighters is 47.9 (±4.6) years old and the average time of service in the Department is 17.4 (±4.2) years. The racial/ethnic make-up of this population in the department is: 50% non-Hispanic White, 21% Asian/Pacific Islander, 17% Hispanic/Latino, and 13% African American, which is reflected by recruited firefighter participants. Among the office workers, the average age is 47 years old and most have worked an average of 14.0 years for the City and County of San Francisco. The racial and ethnic make-up of this workforce was statistically similar to that of the firefighters, with a higher percentage of non-Hispanic Asian/Pacific Islanders (25%).

**Table 1.**
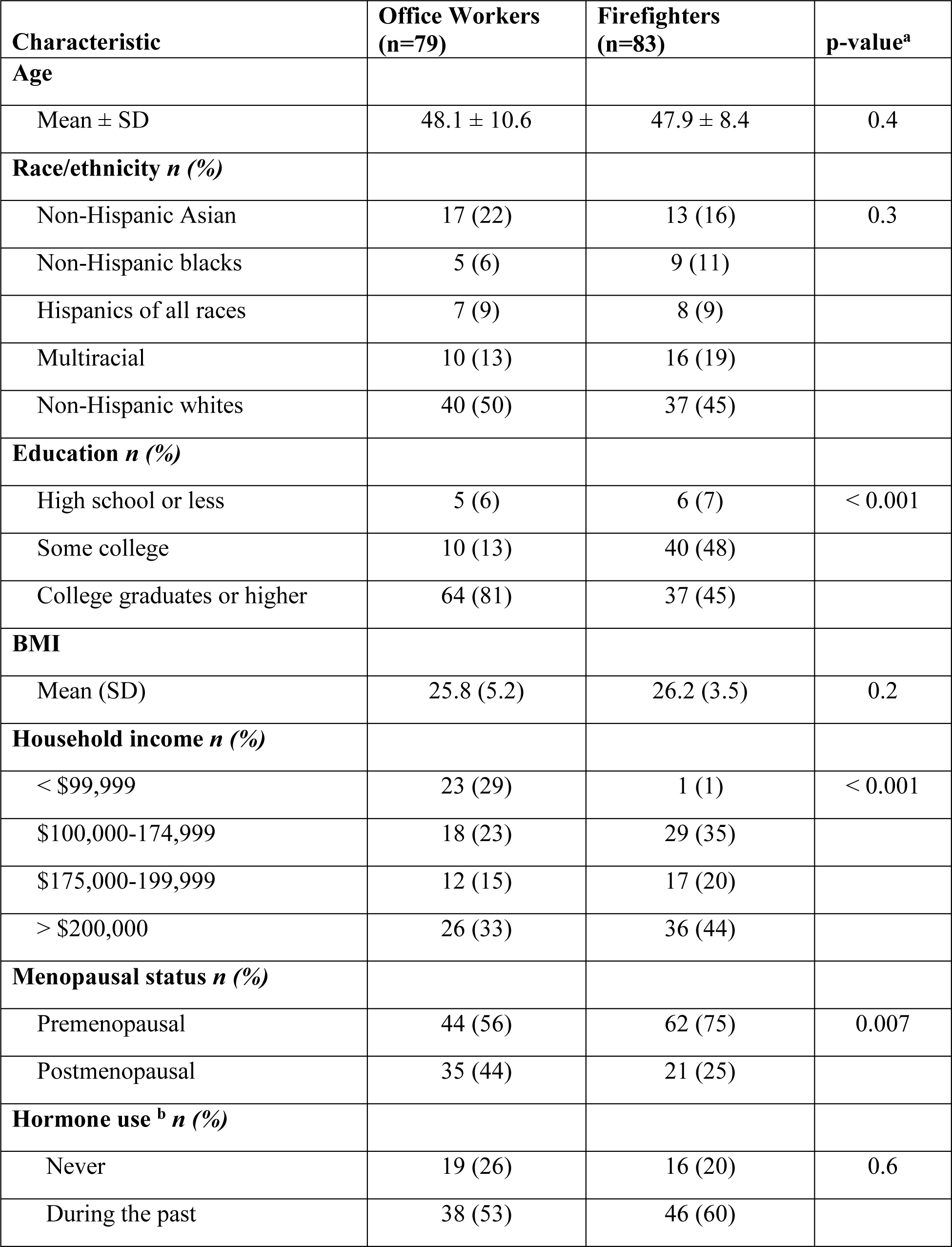
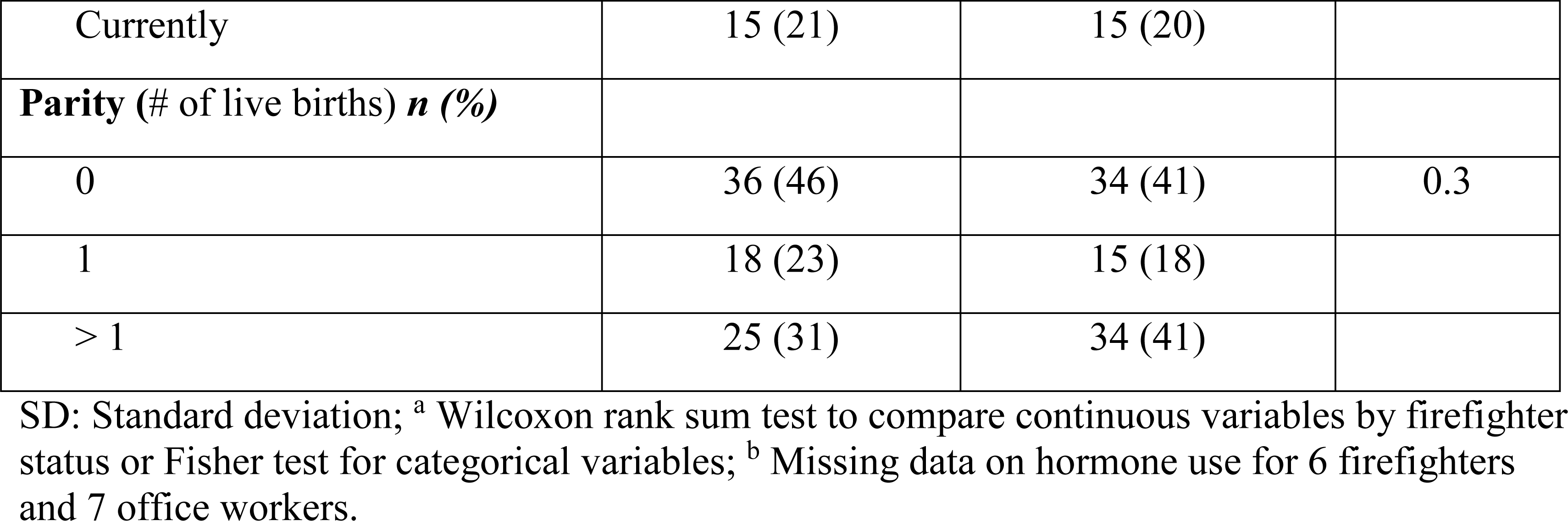
WFBC study population characteristics.

Overall, the firefighters and office workers were similar in terms of average age, race/ethnicity, body mass index (BMI), parity, and hormone use. However, the household income for firefighters was significantly higher when compared to office workers, probably because of the relatively higher compensation rate for firefighting versus office or clerical work. There were significantly more premenopausal women in the firefighter group. Finally, office workers had a higher proportion of college graduates than the firefighters.

### Suspect screening analysis of serum samples

Our general suspect screen analysis identified 12,051 candidate compounds across all serum samples, which were then compared to 740 chemical formulas from the WFBC database. Retention time correction identified 300 chemical formulas, with multiple retention times per formula such that there were 620 putative chemicals in the firefighter and office worker samples. These included phthalate metabolites, phosphate flame retardants (PFRs) and their metabolites, phenols, pesticides, nitro- and nitroso-compounds, and per- and polyfluoroalkyl substances (PFASs). Figure 1 shows the number of chemical suspect hits per participant for each chemical class. A large number of chemicals detected in FF and OW using this analytical method were phenols and phthalate metabolites. The average cumulative number of suspect chemicals detected was 73 (minimum: 45, maximum: 109) and 70 (minimum: 45; maximum: 100) in FF and OW, respectively. Thus, the non-targeted LC-QTOF/MS data acquisition in ESI-was able to detect a wide range of suspect organic acids that include many common commercial chemicals.

**Figure 1:**
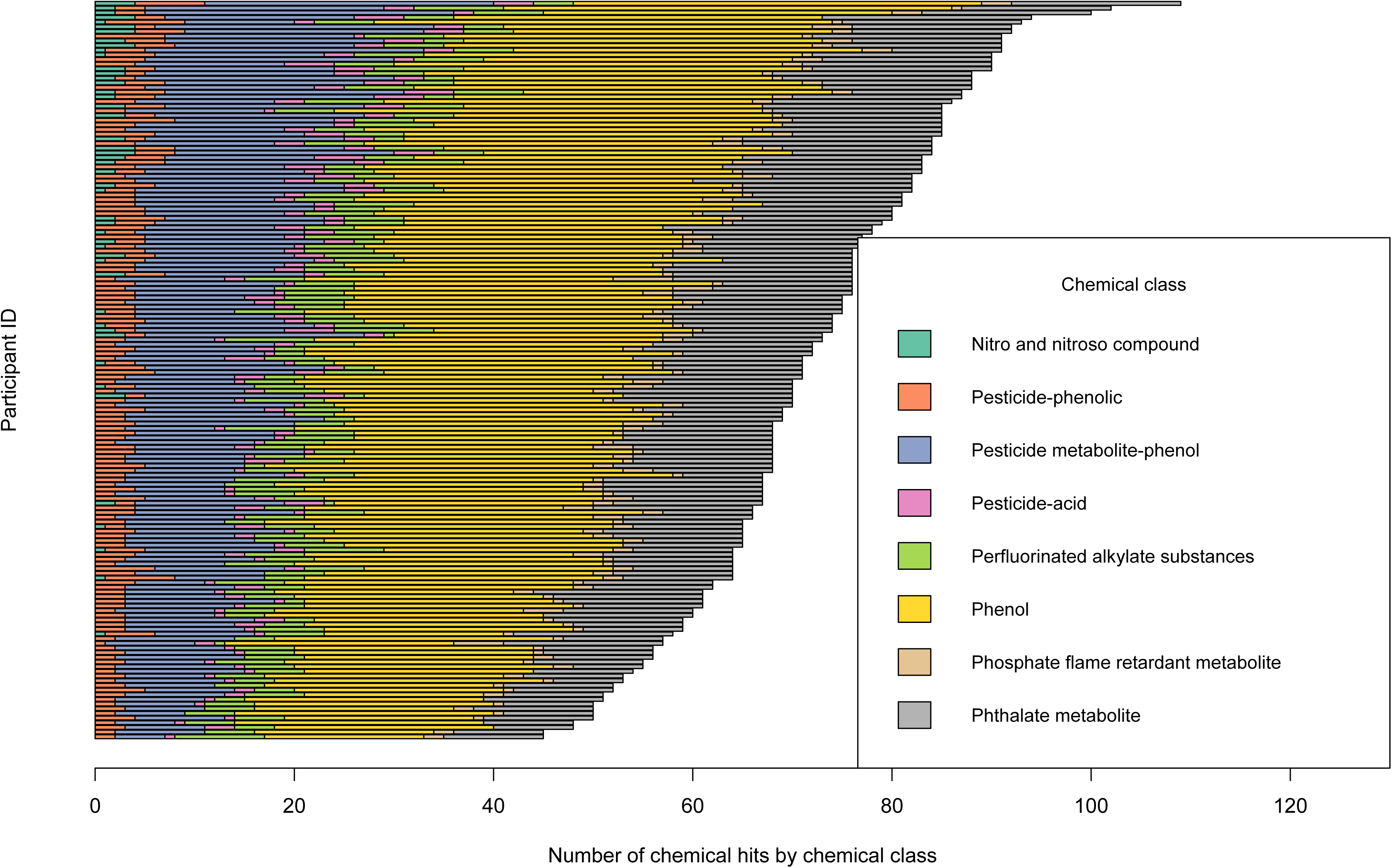
Cumulative number of WFBC database chemicals detected with LC-QTOF/MS ESI-in serum samples from 162 study participants (mean=72; min=45; max=109).

### Chemical restriction and prioritization for validation

We identified 71 chemicals that were: 1) more abundant in firefighters or 2) ubiquitous and not already in NHANES or 3) tagged as a potential concern for breast cancer. Sixty-three of these chemicals satisfied only one criteria, and eight satisfied more than one. We further reduced this list to chemicals that had commercially available authentic standards, leaving 54 to be considered for validation. These chemicals included phenols such as bisphenol F and some alkylphenols, phthalate metabolites, PFAS, flame retardant metabolites, nitroso-compounds, and pesticides (See Table S2). None of the chemicals had significantly different detection frequencies or peak areas in FF versus OW, but many had smaller differences. Fewer than half were identified as mammary carcinogens or developmental disruptors. We scored the remaining 54 chemicals based on indications of toxicity and exposure potential (Figure 2, Table S2).

**Figure 2:**
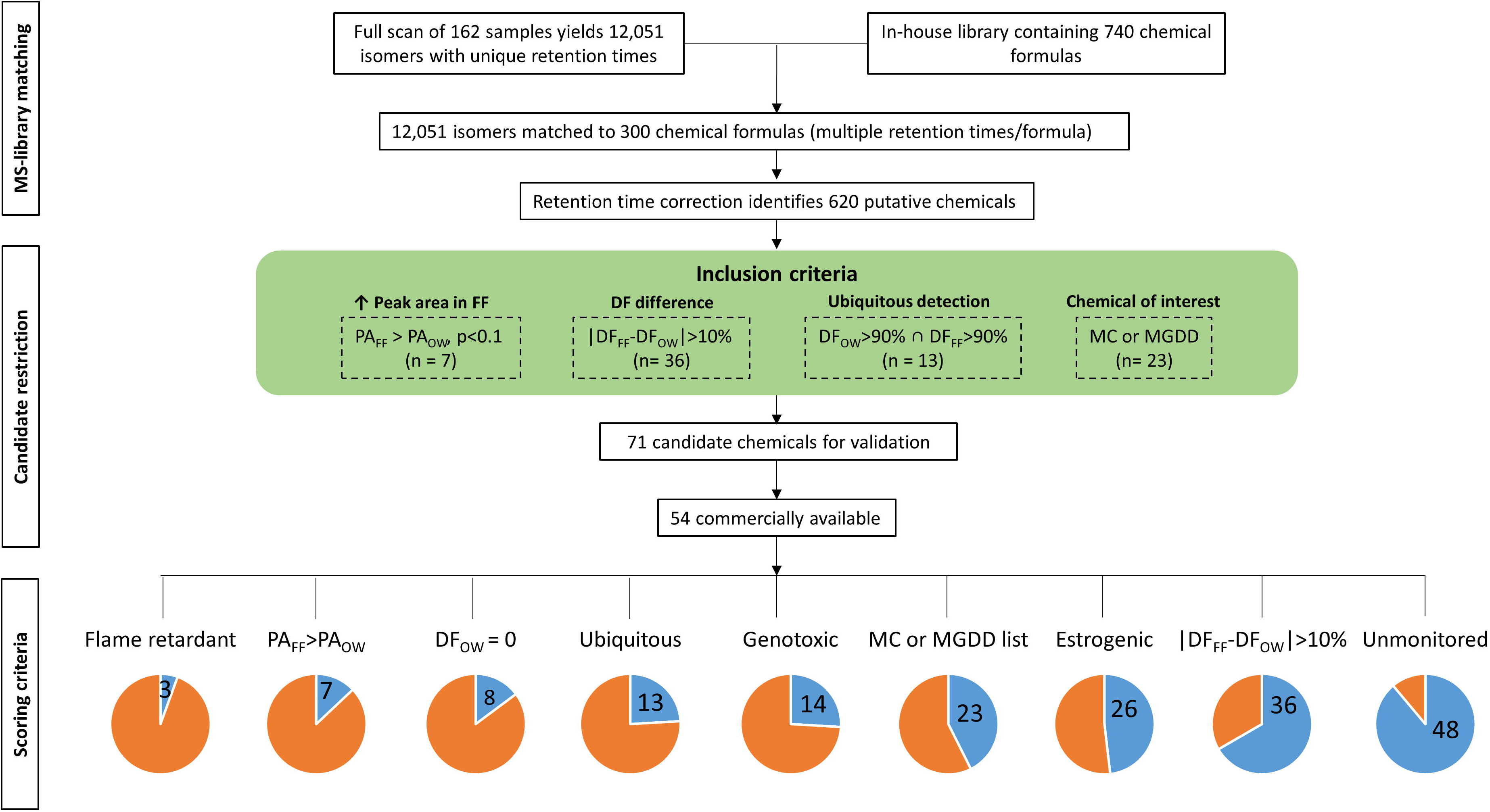
Scoring and ranking of chemicals detected by LC-QTOF. Figure 2 legend: PA= peak area; FF= firefighters; OW = office workers; DF = detection frequency, MC= mammary carcinogen; MGDD = mammary gland developmental disruptor

We selected chemicals for analytical validation after reviewing the priority scores across nine criteria for the 54 chemicals along with data on uses, toxicity and sources (Table S2 provides this information for all 71 candidate chemicals).

Table 2 shows the top 20 scoring candidate chemicals and indicates the priority rank and whether the chemical was included in the confirmation testing. For example, 2,4-bis(1,1-dimethylethyl) phenol had the top ranking, meeting six of the nine criteria (Table 2) and was selected for validation. Three nitro- and nitroso compounds with high scores, including 1-ethylnitroso-3-(2-oxopropyl)-urea, 1-ethylnitroso-3-(2-hydroxyethyl)-urea and 1-amyl-1-nitrosourea were eliminated because although our initial search indicated standards were available, the cost to purchase them was prohibitive. Bis(1,3-dichloro-2-propyl) phosphate (BDCIPP) was excluded because it was already being targeted for analysis in this cohort. Estradiol was excluded because it is endogenous and Nifurdazil, an anti-bacterial agent, was excluded because we were not targeting pharmaceuticals. We included the remaining 14 priority chemicals in the confirmation testing.

**Table 2:** Twenty highest scoring chemicals prioritized for validation

| Chemical name | Class | Rank | DF FF (%) | DF OW (%) | Mean peak area FF | Mean peak area OW | Flame retardant | DF > 90% in FF and OW | \|DF_FF - DF_OW\| > 10% | T-test PA p<0.1 | Unmonitored <sup>a</sup> | Genotoxic | Estrogenic | OW non-detect | MC list | Total Score | Validation status |
| --- | --- | --- | --- | --- | --- | --- | --- | --- | --- | --- | --- | --- | --- | --- | --- | --- | --- |
| 2,4-bis(1,1-dimethylethyl)phenol | Phenol | 1 | 82 (100%) | 76 (100%) | 9.17E+05<sup>†</sup> | 7.66E+05 | 0 | 1 | 0 | 1 | 1 | 1 | 1 | 0 | 0 | 5 | S |
| benzyl p-hydroxybenzoate (PHBB) or <sup>b</sup> | Phenol | 2 | 16 (19.5%) | 6 (7.9%) | 2.98E+04 | 2.12E+04 | 0 | 0 | 1 | 0 | 1 | 1<br>0 | 1 | 0 | 0 |  | S |
| 2-hydroxy-4-methoxybenzophenone -2 (BP-3) |  |  |  |  |  |  |  |  |  |  |  |  |  |  |  |  |  |
| 4-hexyloxyphenol | Phenol | 3 | 81 (98.8%) | 71 (93.4%) | 1.04E+05* | 7.51E+04 | 0 | 1 | 0 | 1 | 1 | 1 | 1 | 0 | 0 | 5 | S |
| benzyl p-hydroxybenzoate (PHBB) or <sup>b</sup> | Phenol | 4 | 30 (36.6%) | 38 (50%) | 6.04E+04 | 9.68E+04 | 0 | 0 | 1 | 0 | 1 | 1<br>0 | 1 | 0 | 0 |  | S |
| 2-hydroxy-4-methoxybenzophenone -2 |  |  |  |  |  |  |  |  |  |  |  |  |  |  |  |  |  |
| bisphenol F | Phenol | 5 | 10 (12.2%) | 0 (0%) | 4.98E+05 | NA | 0 | 0 | 1 | 0 | 1 | 0 | 1 | 1 | 0 | 4 | S |
| 4-butoxyphenol | Phenol | 6 | 77 (93.9%) | 71 (93.4%) | 7.21E+04 | 8.58E+04<sup>†</sup> | 0 | 1 | 0 | 1 | 1 | 0 | 0 | 0 | 0 | 3 | S |
| 2,3,6-trimethylphenol | Phenol | 7 | 18 (22%) | 7 (9.2%) | 2.04E+04 | 1.15E+04 | 0 | 0 | 1 | 0 | 1 | 0 | 0 | 0 | 0 | 2 | S |
| 1-ethylnitroso-3-(2-oxopropyl)-urea | Nitro and Nitroso | 8 | 14 (17.1%) | 10 (13.2%) | 2.54E+04 | 2.09E+04 | 0 | 0 | 0 | 0 | 1 | 0 | 0 | 0 | 1 | 2 | E-No std |

|  | Compound |  |  |  |  |  |  |  |  |  |  |  |  |  |  |  |  |
| --- | --- | --- | --- | --- | --- | --- | --- | --- | --- | --- | --- | --- | --- | --- | --- | --- | --- |
| perfluorooctanesulfona midoacetate (PFOSAA) | PFAS | 9 | 16<br>(19.5%) | 25<br>(32.9%) | 3.94E+0<br>4 | 4.56E+0<br>4† | 0 | 0 | 1 | 1 | 1 | 0 | 0 | 0 | 0 | 3 | S |
| diphenyl phosphate (DPP) | Phosphate<br>Flame<br>Retardant<br>metabolite | 10 | 45<br>(54.9%) | 39<br>(51.3%) | 1.57E+0<br>4 | 1.68E+0<br>4 | 1 | 0 | 0 | 0 | 0 | 0 | 0 | 0 | 0 | 1 | S |
| bis(1,3-dichloro-2-propyl) phosphate (BDCIPP) | Phosphate<br>Flame<br>Retardant<br>metabolite | 11 | 2<br>(2.4%) | 1<br>(1.3%) | 1.35E+0<br>4 | 1.13E+0<br>4 | 1 | 0 | 0 | 0 | 0 | 0 | 0 | 0 | 1 | 2 | E- target<br>analyte |
| 4-phenethylphenol | Phenol | 12 | 82<br>(100%) | 76<br>(100%) | 1.35E+0<br>5 | 1.43E+0<br>5† | 0 | 1 | 0 | 1 | 1 | 0 | 1 | 0 | 0 | 4 | S |
| 4-heptyloxyphenol <sup>b</sup><br>(isomer 1) | Phenol | 13 | 31<br>(37.8%) | 21<br>(27.6%) | 6.60E+0<br>4 | 6.87E+0<br>4 | 0 | 0 | 1 | 0 | 1 | 0 | 1 | 0 | 0 | 3 | S |
| Nifurdazil | Nitro and<br>Nitroso<br>Compound | 14 | 4<br>(4.9%) | 3<br>(3.9%) | 2.37E+0<br>4 | 1.07E+0<br>4 | 0 | 0 | 0 | 0 | 1 | 1 | 0 | 0 | 1 | 3 | E -<br>medication |
| 4-heptyloxyphenol <sup>b</sup><br>(isomer 2) | Phenol | 15 | 51<br>(62.2%) | 55<br>(72.4%) | 2.89E+0<br>5 | 2.55E+0<br>5 | 0 | 0 | 1 | 0 | 1 | 0 | 1 | 0 | 0 | 3 | S |
| 1-ethylnitroso-3-(2-hydroxyethyl)-urea | Nitro and<br>Nitroso<br>Compound | 16 | 3<br>(3.7%) | 2<br>(2.6%) | 1.57E+0<br>4 | 1.57E+0<br>4 | 0 | 0 | 0 | 0 | 1 | 0 | 0 | 0 | 1 | 2 | E-No std |
| 1-amyl-1- nitrosourea | Nitro and<br>Nitroso<br>Compound | 17 | 7<br>(8.5%) | 11<br>(14.5%) | 3.56E+0<br>4 | 2.33E+0<br>4 | 0 | 0 | 0 | 0 | 1 | 0 | 0 | 0 | 1 | 2 | E-No std |
| ethyl-p-hydroxybenzoate (ethyl paraben) | Phenol | 18 | 52<br>(63.4%) | 35<br>(46.1%) | 1.10E+0<br>5 | 1.57E+0<br>5* | 0 | 0 | 1 | 1 | 0 | 0 | 1 | 0 | 0 | 3 | S |
| 1-allyl-1-nitrosourea | Nitro and<br>Nitroso<br>Compound | 19 | 12<br>(14.6%) | 5<br>(6.6%) | 7.25E+0<br>4 | 3.96E+0<br>4 | 0 | 0 | 0 | 0 | 1 | 0 | 0 | 0 | 1 | 2 | S |
| estradiol | Steroid | 20 | 1<br>(1.2%) | 0<br>(0%) | 1.03E+0<br>4 | NA | 0 | 0 | 0 | 0 | 0 | 1 | 1 | 1 | 1 | 4 | E-endogenous |

<sup>a</sup> Unmonitored in NHANES or Biomonitoring California; <sup>b</sup> these are isomers and could not be distinguished based on molecular mass;
\*p<0.1; † p<0.05; FF = firefighter; OW = office worker; DF = detection frequency; PA = peak area; RT=retention time;
MC=mammary carcinogen; E = eliminated for validation; S = selected for validation; LOD = limit of detection; std=standard

### Validation

Authentic standards of the 14 selected chemicals were analyzed by LC-QTOF/MS to evaluate their match with retention times and mass spectra in the samples. Retention times for chemical candidates and authentic standards, exact masses, and validation status are listed in Table 3. Eight chemicals were validated, including: 2,4-bis(1,1-dimethylethyl)phenol, 2-hydroxy-4-methoxybenzophenone −2, bisphenol F, perfluorooctanesulfon-amidoacetate (PFOSAA), diphenyl phosphate (DPP), ethyl-p-hydroxybenzoate (ethyl paraben), benzyl p-hydroxybenzoate (PHBB), and 4-hexyloxyphenol.

**Table 3:** Retention time and exact mass for chemicals selected for validation

| Chemical name | Chemical class | # of isomers | Mean RT for serum samples | RT lab standard | Validation status |
| --- | --- | --- | --- | --- | --- |
| 2,4-bis(1,1-dimethylethyl) phenol | Phenol | 4 | 4.33, 5.25, 5.48, 6.73 | 6.72 | ✓ |
| 2-hydroxy-4-methoxybenzophenone (BP-3)) | Phenol | 2 | 4.33, 5.25 | 5.30 | ✓ |
| bisphenol F | Phenol | 2 | 3.91 | 4.00 | ✓ |
| perfluorooctanesulfonamidoacetate (PFOSAA) | PFC | 1 | 5.93 | 5.95 | ✓ |
| diphenyl phosphate (DPP) | Phosphate Flame Retardant metabolite | 1 | 3.86 | 3.90 | ✓ |
| ethyl-p-hydroxybenzoate (ethyl paraben) | Phenol | 2 | 2.21, 3.80 | 2.30 | ✓ |
| benzyl p-hydroxybenzoate (PHBB) | Phenol | 2 | 4.33, 5.25 | 4.40 | ✓ |
| 4-hexyloxyphenol<sup>1</sup> | Phenol | 1 | 5.81 | 5.80 | ✓<sup>a</sup> |
| 4-butoxyphenol | Phenol | 1 | 4.19 | 5.10 | ✗<sup>b</sup> |
| 2,3,6-trimethylphenol | Phenol | 2 | 3.97 | 4.25 | ✗<sup>b</sup> |
| 4-phenethylphenol | Phenol | 1 | 5.71 | 6.02 | ✗<sup>b</sup> |
| 4-heptyloxyphenol (2 isomers) | Phenol | 1 | 5.09 | 6.22 | ✗<sup>b</sup> |
| 1-allyl-1-nitrosourea | Nitro and Nitroso compound | 1 | 0.76 | 1.20 | ✗<sup>b</sup> |

<sup>a</sup> validated but with high LOD, <sup>b</sup> not validated because of retention time mismatch

We found that retention times in participants’ serum did not match those of the standards for six chemicals: 1-allyl-1-nitrosourea, 4-butoxyphenol, 2,3,6-trimethylphenol, 4-phenethylphenol, and two isomers for 4-heptyloxyphenol.

## DISCUSSION

The goal of this study was to apply a general suspect screening approach to identify novel exposures to previously understudied chemicals – of particular relevance to firefighting and breast cancer etiology -- among a cohort of women firefighters compared to office worker controls. We used LC-QTOF/MS to screen for the presence of 740 chemicals of interest in serum from women firefighters and office workers. Accurate masses of chemical suspects were tentatively matched with exact masses from the WFBC chemical database developed for this study; chemical suspects were then prioritized for validation based on criteria related to exposure profiles between the two groups as well as toxicity information, expected exposure patterns, and whether they are currently biomonitored in major surveillance programs or not.

We detected 620 chemicals that matched 300 different molecular formulas, including phthalate metabolites, phosphate flame retardants and their metabolites, phenols, pesticides, nitro- and nitroso-compounds, and PFAS in both FF and OW. The average number of suspect chemicals detected was 73 and 70 in FF and OW, respectively. Eight of the 20 prioritized chemicals were validated by analysis with a known standard and will ultimately be quantified in the samples. This approach presents a novel and powerful method for using suspect screening in a cohort of female firefighters to reveal exposures to previously unstudied chemicals and to prioritize compounds for confirmation.

Among the eight chemicals whose identity was validated by matching retention time and MS/MS fragmentation of a known standard, the results suggested that exposures were different between firefighters and office workers for most of them, although the magnitude of the differences was modest. Based on statistically significant differences in peak area, firefighters had higher relative levels of exposure for 2,4-bis(1,1-dimethylethyl) phenol, and office workers for PFOSAA and ethyl paraben (Table 2). Firefighters appeared to have slightly higher detection frequencies for 2-hydroxy-4-methoxybenzophenone (BP-3), bisphenol F, PFOSAA and ethyl paraben, and office workers had a higher detection frequency for PHBB.

The validated chemicals included two phenols, (bisphenol F and PHBB), which are used as bisphenol-A substitutes (Ng et al. 2015), and BP-3, which is a UV filter in sunscreens, textiles, and other products. The chemical 2,4-bis(1,1-dimethylethyl) phenol (aka 2,4-di-tert butyl phenol), is listed as a manufacturing chemical and a fuel additive, however since it was detected in all of the participants it may have some common consumer use or be a metabolite of a common exposure (CID 7311) (Kim et al. 2016). It is interesting to note the similarity to 4-tert butyl phenol—a stronger estrogen mimic that is ubiquitous in residential settings (Rudel et al. 2003). Ethyl paraben is an antifungal preservative found in cosmetics, toys, sunscreen and pesticides (Guo and Kannan 2013). A PFAS chemical, PFOSAA, was also validated. Previous studies have reported higher firefighting exposures for PFASs (Laitinen et al. 2014; Rotander et al. 2015), and findings of targeted analysis for PFASs in this cohort are forthcoming (Trowbridge et al. in prep). Originally a metabolite of an active ingredient in Scotchgard stain and water repellant, PFOSAA is listed as an automotive, construction-related and cleaning chemical, as well as an inert pesticide ingredient (CID 23691014) (Kim et al. 2016). It may also be found in firefighting foams. Diphenyl phosphate, a common metabolite of the flame retardant and plasticizer triphenyl phosphate (Cooper et al. 2011), appeared to have similar concentrations in firefighters and office workers.

Among the few studies previously conducted on firefighters, one (Waldman et al. 2016) observed higher exposures to environmental phenols (i.e. bisphenol A, triclosan, benzophenone-3 and methyl paraben) among Southern California firefighters compared to the general population. Since this study also investigated firefighters from California, it is difficult to decipher whether the prevalent exposures to phenols are specifically related to firefighting activities or simply more prevalent among California populations in general.

The phenols and PFAS chemicals that were validated in this study have estrogenic activity (Table 2) or are of concern for a diverse set of toxicity endpoints, such as effects on kidney, liver, lipid metabolism, growth and development, mammary gland development, and immunotoxicity (Post et al. 2017). While there were tentative matches to nitro and nitroso chemicals, which are of interest because of their genotoxicity and carcinogenicity (Table 2), we were not able to validate any of these compounds, either because the retention time did not match the known standard or we could not obtain the standard.

The success of this general suspect screening technique to identify novel chemical exposures in environmental and occupational health studies could be improved further if there were chemical databases that contain mass spectral information about diverse chemicals of interest. Because most public metabolomics databases, such as HMDB, Metlin or T3DB, contain few entries for environmental chemicals (e.g. HMDB contains 163 entries for toxin/pollutant) and there are no extensive mass spectral databases of environmental chemicals currently available, we instead made comparisons to 740 chemicals in our database based on matching exact masses. This approach allowed us to tentatively identify exposures of interest by focusing the search on a set of chemicals of interest and for which the analytical method was optimized. We also demonstrated that this approach can be effective in measuring low abundant chemicals in human serum. For example, PFOS detected using the GSS (Table S2) was also confirmed and quantified using targeted LC-MS/MS (median serum concentrations for the whole cohort were 4.1 ng/mL for PFOS) (Trowbridge et al. in prep).

We were also interested in identifying exposures associated with work practices that are not related to fire events, such as diesel fuel and exhaust from trucks and equipment in the station, flame retardants and PFAS chemicals from firefighting foam and protective gear, chemicals used to clean and gear, and possibly others. Some of the chemicals selected for targeted analyses may be related to workplace exposures such as these, and this suspect screening approach is one way to generate hypotheses about exposures and to prioritize novel compounds for confirmation and quantification using targeted methods.

Our study has several limitations. The sample size is relatively modest, and a larger cohort would have provided more power to detect candidate chemicals that differed between firefighters and office workers. In addition, since most of chemicals we detected are non-persistent, we can expect large intra-individual variability in serum due to temporal variation in exposure. Also, only 15 firefighters had their blood sample collected within 24 hours of working at a fire event, so it may be that the chemicals we detected were not necessarily associated with firefighting activities. One way to better characterize chemicals originating from fighting fires would be to perform a longitudinal analysis in which biospecimens would be collected before and after a fire event (within 12-24h).

Our WFBC general suspect chemical database (740 chemicals) contained only a small fraction of the chemicals that could be important exposures for firefighters and office workers and so we may have missed some important compounds for this study population. The use of larger chemical databases such as the EPA Distributed Structure-Searchable Toxicity (DSSTOX; ~9,000 chemicals) (Richard and Williams 2002) or PubChem (~3,000 chemicals) (Kim et al. 2016) would provide detection of a larger set of chemical suspects. However, increasing the number of chemicals in a general suspect database would likely also increase the number of “hits” (tentative chemical RT matches), making it more challenging to confirm matches and increasing the rate of false positives. Even with our database of 740 chemicals, six, two of which are isomers, of the top 20 tentative chemical matches that we selected for validation showed a retention time (RT) mismatch such that the study serum sample RT did not match the RT generated from a reference standard. Combining LC-QTOF/MS data - collected using a data-independent acquisition approach (i.e. MS/MS fragmentation of as many metabolites as possible in a single acquisition) – with bioinformatics tools such as retention time prediction, in silico MS/MS prediction and molecular networking analysis (Allard et al. 2016; Bessonneau et al. 2017) would help to address this issue. In addition, a careful validation of the chemical identity using an authentic standard is required to avoid reporting false positive matches. Likewise, the number of matching fragmentation peaks required to minimize false positives can be investigated in future studies. Ultimately, the MS/MS spectra generated for any compound provide structural information specific to a compound. This data becomes very valuable for distinguishing isomeric compounds that may have very close retention times in chromatography.

Another limitation is that use of LC/QTOF-MS in negative ionization mode limited the types of chemicals that could be detected to organic acids. The use of complementary platforms such as LC-QTOF/MS in positive ionization mode or GC combined with high resolution MS would expand the investigation to more diverse classes of chemicals. For example, Greer Wallace et al. (Geer Wallace et al. 2017) identified several VOCs and PAHs in firefighters exposed to controlled structure burns using targeted and non-targeted GC-MS analysis of exhaled breath condensate. Some of these chemicals such as benzaldehyde and dimethyl sulfide have been previously associated with smoke/fire and combustion sources while methyl tert-butyl ether is commonly used as an additive to gasoline. Finally, some of the nitroso compounds with high priority scores in our analysis such as 1-amyl-1-nitrosourea and 1-allyl-1-nitrosourea could not be validated because standards were not available.

In summary, we present a general suspect screening approach based on LC-QTOF/MS that can be used to identify novel chemical exposures (i.e. not previously biomonitored) in a way that is not as strictly limited by *a priori* hypotheses required by targeted methods. The approach we used to select chemicals for confirmation integrates information from the serum samples, toxicity and usage databases and expert knowledge to direct attention to chemicals relevant to the health of women firefighters, an understudied yet vulnerable occupational group. Follow-up studies should include targeted analyses to confirm and quantify the identified chemicals in the cohort, identification of potential sources of the exposures, extension of the approach to cover a broader and more diverse chemical space, and assessment of potential associations with health outcomes for validated chemicals.

## Acknowledgements

The authors thank all of the WFBC study participants for their contribution to the study. This work is supported by the California Breast Cancer Research Program #19BB-2900 (RG, VB, RRG, JT, TL, HB, RAR, RMF), the National Institute of Environmental Health Sciences R01ES027051 (AW, RMF) and the San Francisco Firefighter Cancer Prevention Foundation (HB). We thank Anthony Stefani, Emily O’Rourke, Nancy Carmona, Karen Kerr, Julie Mau, Natasha Parks, Lisa Holdcroft, SF Fire Chief Joanne Hayes-White, Incoming SF Fire Chief Jeanine Nicholson, Sharyle Patton, Connie Engel and Nancy Buermeyer for their contributions to the study.

RAR, RG, and VB, are employed at the Silent Spring Institute, a scientific research organization dedicated to studying environmental factors in women’s health. The Institute is a 501(c)3 public charity funded by federal grants and contracts, foundation grants, and private donations, including from breast cancer organizations. HB is former president and member of United Fire Service Women, a 501(c)3 public charity dedicated to supporting the welfare of women in the San Francisco Fire Department.

The authors declare they have no actual or potential competing financial interests.

